# A Sparse Modal Representation for Dolphin Echolocation Transients

**DOI:** 10.64898/2026.10.14.759144

**Authors:** F. P. L. Strijbos

## Abstract

Characterising echolocation provides a non-invasive means of monitoring cetacean populations, health, and behaviour, offering critical data to inform conservation policy and mitigate the impacts of anthropogenic ocean noise. Echolocation clicks are short broadband transients whose internal oscillatory structure is difficult to compare in a sequence using conventional time- or frequency-domain representations. We introduce a Sparse Modal Representation (SMR) in which each click is described as a sparse superposition of delayed damped resonant modes. The representation separates resonant behaviour from temporal localisation, allowing both narrowband and broadband click components to emerge from the same set of resonant building blocks. The methodology combines Burg autoregressive pole estimation, sparse approximation in an over-complete dictionary, and nonlinear parameter refinement to obtain compact and interpretable modal descriptions of individual transients. The method is evaluated using recordings of free-ranging common dolphins (Delphinus delphis) from the DOLPHINFREE project in the Northern Bay of Biscay.

The results demonstrate accurate reconstruction using a limited set of parameters that are repeatable across successive clicks and robust under realistic recording conditions. These conditions include wave, vessel, and sonar interference. Repeatability and robustness are essential for using this parameter set to detect changes in animal behaviour.

## 1 INTRODUCTION

Echolocation clicks of marine mammals have been the subject of extensive research since the discovery of dolphin biosonar (Au, 2015). Dolphins and other toothed whales use echolocation signals for navigation, prey detection, and sensing their direct environment. Research by Leighton et al. (2004) suggests dolphins are even able to counteract the effects of attenuation in bubbly water. Echolocation signals are commonly described as broadband impulsive clicks. These are typically characterised using temporal and spectral descriptors such as Inter-Click Interval (ICI), amplitude, dominant frequency, bandwidth, and quality factor (Wahlberg et al., 2011). These descriptors effectively detect and analyse behaviour, but they only provide aggregate measures of transient phenomena and offer limited insight into waveform-generation mechanisms. Consequently, several authors have proposed explicit waveform models for dolphin clicks. The pioneering work of Kamminga (Kamminga and Wiersma, 1981; Kamminga et al., 1996) demonstrated that toothed whale clicks can be represented as the sum of one or more Gabor functions. In this interpretation, the first Gabor function represents the emitted source signal, while the additional Gabor functions account for internal reflections within the sound-production system. More recently, Wei et al. (2018, 2024) employed both a narrowband damped sinusoid and a broadband, 90° phase-shifted Ricker wavelet as alternative source functions in finite-element simulations to reproduce measured dolphin clicks. Rather than using these parametric models or non-parametric Fourier or wavelet representations, the proposed representation allows both narrowband and broadband click structures to emerge naturally from the sparse superposition of delayed damped resonances. This builds upon mode-superposition methods used, for example, in earthquake engineering (Borino and Muscolino, 1986).

Our method provides a Sparse Modal Representation (SMR) that identifies the dominant resonances and their temporal activations within individual dolphin echolocation clicks. Because the representation describes the observed waveform rather than assuming a specific source mechanism, it remains compatible with several existing physical interpretations of dolphin click production, including models based on internal reflections (Kamminga et al., 1996), while providing a common framework in which narrowband and broadband waveform components can be analysed simultaneously.

## 2 METHODOLOGY OVERVIEW

The proposed methodology is introduced using a synthetic signal made by combining three damped resonances with prescribed frequencies and quality factors. This synthetic signal is repeated three times to mimic a typical click sequence with a 3 ms interclick interval.

Because these modal parameters are known, this example illustrates how the method can recover the constituent resonances while introducing the sparse modal representation used throughout the remainder of the paper. The same visualisation is subsequently applied to dolphin echolocation transients in Section 5.

Figure 1(c) shows the synthetic transient in the compressed time scale used throughout this paper to place individual transients within a sequence of clicks. When any preprocessing is applied, both the raw and filtered waveforms are displayed. Individual transients are identified using a peak-picking algorithm. Each transient is displayed using a time axis expanded from the detected onset. The analysis window is indicated by the horizontal range between Figure 1(a) and (b).

**FIGURE 1.**
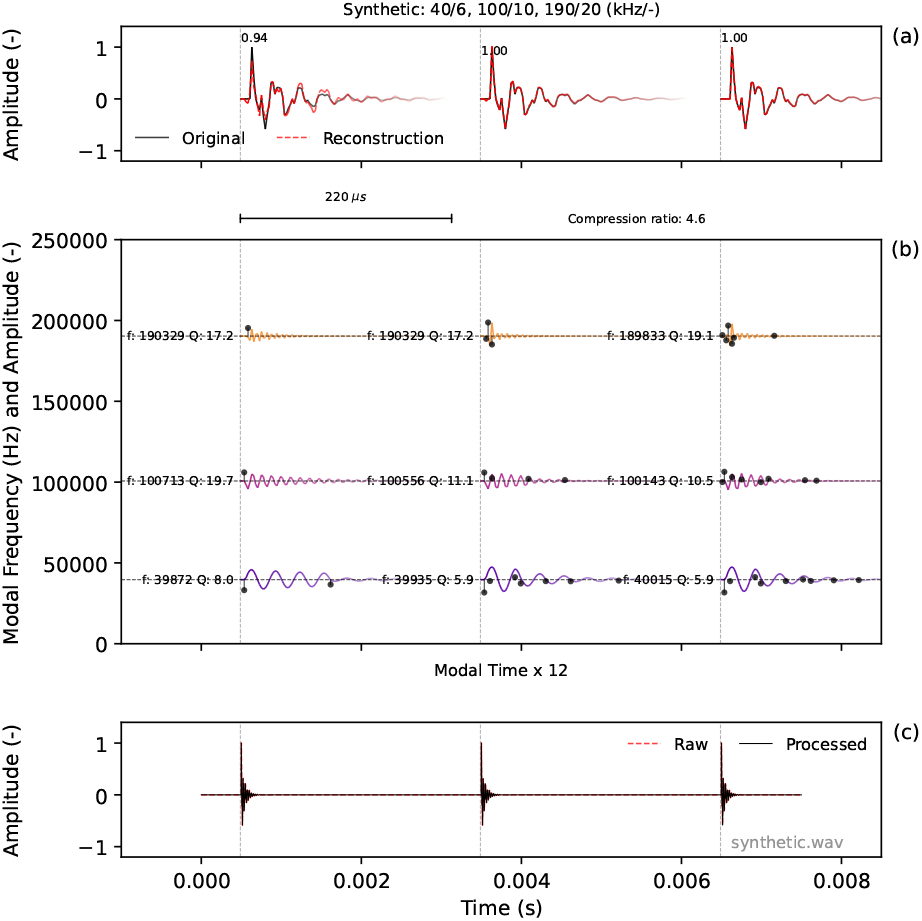
Sparse modal representation of a synthetic transient repeated three times to form a click sequence with a 3 ms interclick interval. The transient is made by adding three damped resonances (frequency/quality factor: 40 kHz/6, 100 kHz/10 and 190 kHz/20). The dictionary is built from three estimated damped resonances, using, from left to right, 4, 14, or 24 atoms.

The transients are first analysed to estimate their dominant resonant frequencies and damping rates. These resonances define the principal oscillatory modes contributing to the recorded waveform. For each resonance, a family of delayed, localised damped oscillations is generated, forming a dictionary of elementary building blocks (atoms). A sparse optimisation procedure then selects only a small subset of these atoms to reconstruct the transient while minimising model complexity. The selected atoms are then grouped by parent resonance, such that a limited number of modal responses describe the temporal contribution of each resonant mode. The compression ratio, which is the ratio of the number of samples in the analysis window to the number of atoms required to represent the transient, measures sparsity.

Figure 1(a) compares the original transient with its reconstruction obtained by summing the grouped modal responses using (left to right) 4, 14, or 24 atoms. The value above the waveform denotes the correlation coefficient between the original and reconstructed signals, which measures reconstruction quality. With just 4 atoms, a compression by a factor of 27.5 is made, and a correlation coefficient of 0.96. With such a low number of atoms, the representation stays close to the derived parametric Autoregressive model. When we use 24 atoms, the correlation coefficient increases to 1, but the damping ratios are within 1% to 23% of the theoretical value. Increasing the reconstruction to 24 atoms reduces this error to less than 5%, but the reconstruction is more akin to a non-parametric method like a wavelet or Fourier reconstruction.

Figure 1(b) presents the sparse modal representation. Each horizontal baseline corresponds to one resonant mode and is positioned at its estimated frequency. The coloured traces show the temporal evolution of the grouped modal responses, while the black stems indicate the amplitudes and temporal locations of the atoms selected by the sparse optimisation.

The annotated frequency and quality factor are refined by fitting a localised damped resonance directly to each grouped modal response. This refinement improves the estimates of resonant frequency, damping, and amplitude while providing confidence measures based on the uncertainty of the fitted parameters. Modes whose refined parameters do not satisfy the prescribed confidence criteria are excluded from the reported modal representation. In practice, these rejected modes contribute negligibly to the reconstructed waveform, so the reconstruction is essentially unaffected.

Because the modal parameters of the synthetic transient are known exactly, the example demonstrates that, under idealised conditions, resonant frequencies and quality factors are recovered to within approximately 1% and 5%, respectively.

In Figure 2, for each repeating synthetic transient, one, two, or three resonant frequencies are estimated, and a dictionary is constructed using 40 atoms. This high number of atoms accurately reconstructs the signal even when only one of the three resonances is used to build the dictionary. Such a reconstruction is more akin to non-parametric methods, though the atoms don’t form a complete basis like a wavelet transform.

**FIGURE 2.**
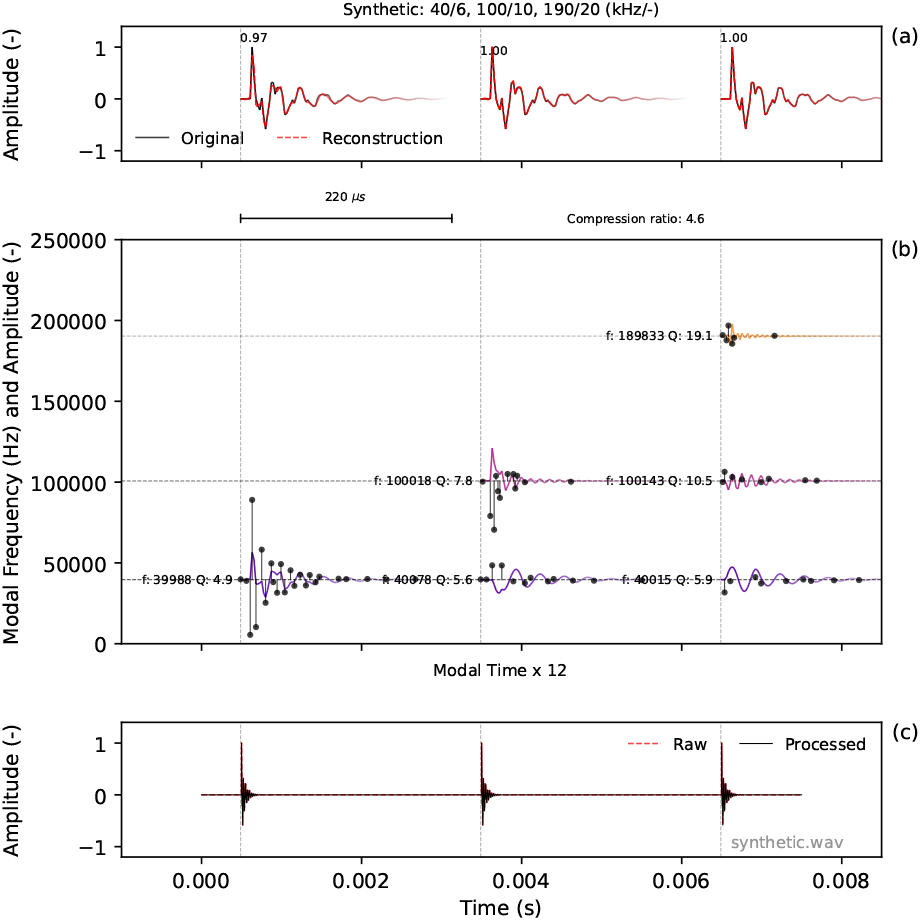
Sparse modal representation of the same synthetic as in Figure 1, but using a dictionary built with 24 atoms created from one, two, or three estimated damped resonances.

We use the modal representation illustrated using these synthetics to investigate the repeatability and robustness of the estimated modal parameters for dolphin echolocation transients.

## 3 DETAILED METHOD

Figure 3 provides an overview of the processing stages comprising the proposed Sparse Modal Decomposition. A detailed mathematical breakdown is given in the appendix. The individual steps are described below in the order shown in the flowchart.

**FIGURE 3.**
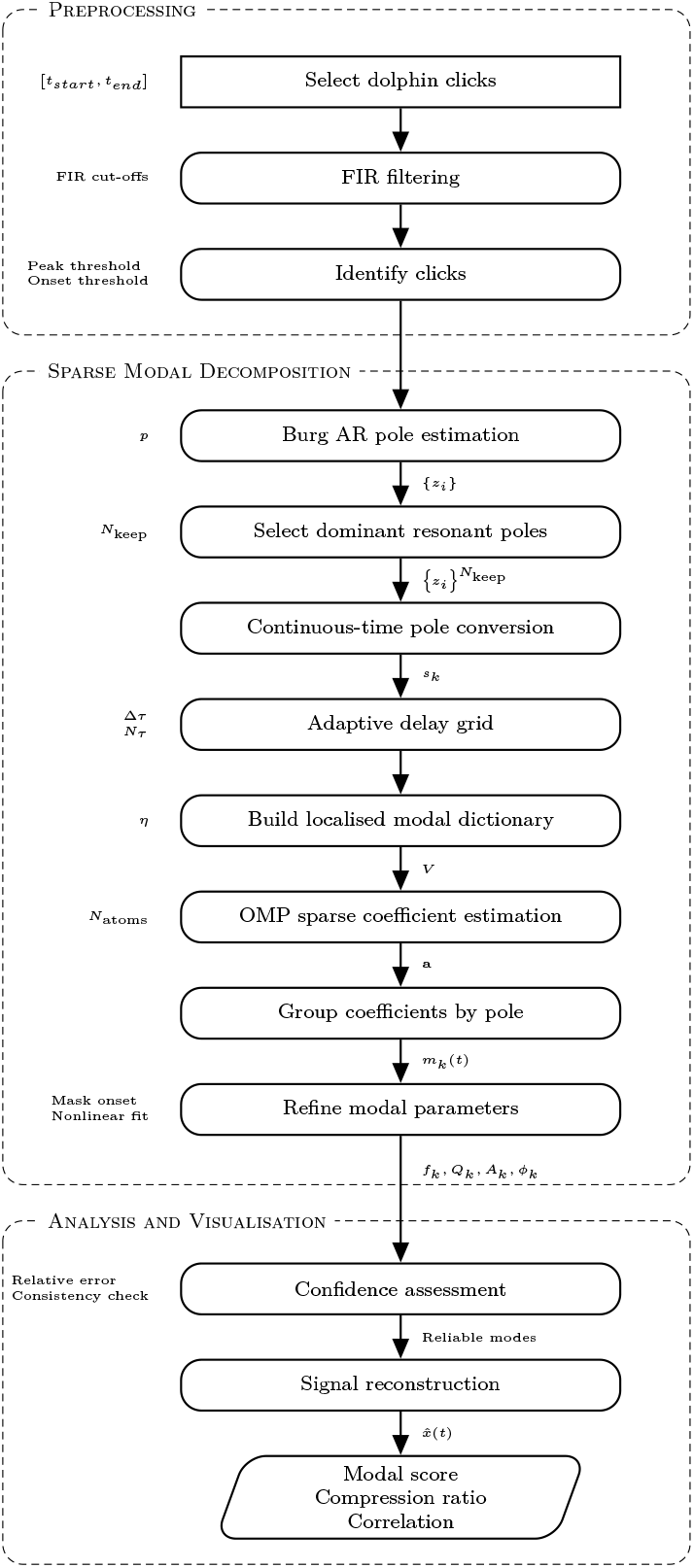
Flowchart summarising the processing stages of the proposed Sparse Modal Decomposition.

The first step of the proposed Sparse Modal Decomposition is the selection of a time interval containing one or more transient events. Processing multiple consecutive transients within a single analysis window enables comparison of the resulting modal representations while preserving their temporal relationship within the recording.

Optional preprocessing is then applied to suppress unwanted frequency components. Depending on the recording conditions, zero-phase Finite Impulse Response (FIR) high-pass, low-pass, or band-pass filtering may be employed to reduce low-frequency wave- and boat noise or narrowband anthropogenic interference (e.g., from an echo-sounder). Zero-phase filtering is used throughout to avoid introducing artificial phase shifts or timing offsets that could bias the subsequent onset detection and modal decomposition.

Local maxima were identified using a peak detection algorithm based on local amplitude maxima exceeding a prescribed threshold. The onset of each transient was determined by searching backwards from the detected peak until the signal amplitude fell below a prescribed fraction of the peak amplitude. A fixed, specified time interval then defines the transient window. The sparse modal decomposition begins by estimating the poles of an autoregressive (AR) model. In the present work, the initial pole estimates are obtained using Burg’s method (Burg, 1975), a parametric AR estimation technique with good frequency resolution for short data records and the ability to resolve closely spaced resonances. These pole estimates provide the initial modal frequencies and damping rates from which the sparse modal dictionary is constructed.

Burg’s autoregressive coefficients were estimated using an in-house implementation of Burg’s algorithm. Because the analysed signals consist of isolated transients rather than stationary time series, a smooth error-function weighting was applied to the prediction error, progressively increasing the contribution of samples following the transient onset. This weighting improves the estimation of the dominant resonant poles while reducing sensitivity to pre-onset noise.

From the estimated poles, only the *N*_*keep*_ dominant resonant pole pairs are retained. This reduces the risk of overfitting the relatively short analysis window. At the same time it exploits the observation that a small number of dominant modes can describe many resonant systems. The retained poles provide the basis for constructing the sparse modal dictionary.

The poles estimated by Burg’s algorithm are discrete-time quantities and therefore depend on the sampling interval. To construct a dictionary of delayed modal atoms with physically meaningful, sampling-independent frequencies and damping rates, and to permit arbitrary delay locations, the discrete poles *z*_*i*_ are mapped to the continuous-time s-plane using

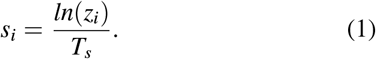

Rather than relying on highly damped poles to reproduce broadband transients, the proposed representation exploits interference between delayed atoms associated with the same lightly damped resonance. By varying the relative delay and coefficient signs, the effective temporal support of the modal response can be adjusted independently of its resonant frequency and damping. As illustrated in Figure 4, neighbouring atoms separated by approximately one half-period can combine through constructive and destructive interference, producing highly localised transient responses while preserving the underlying resonant frequency. This enables short-duration broadband features to be represented without introducing additional highly damped poles.

**FIGURE 4.**
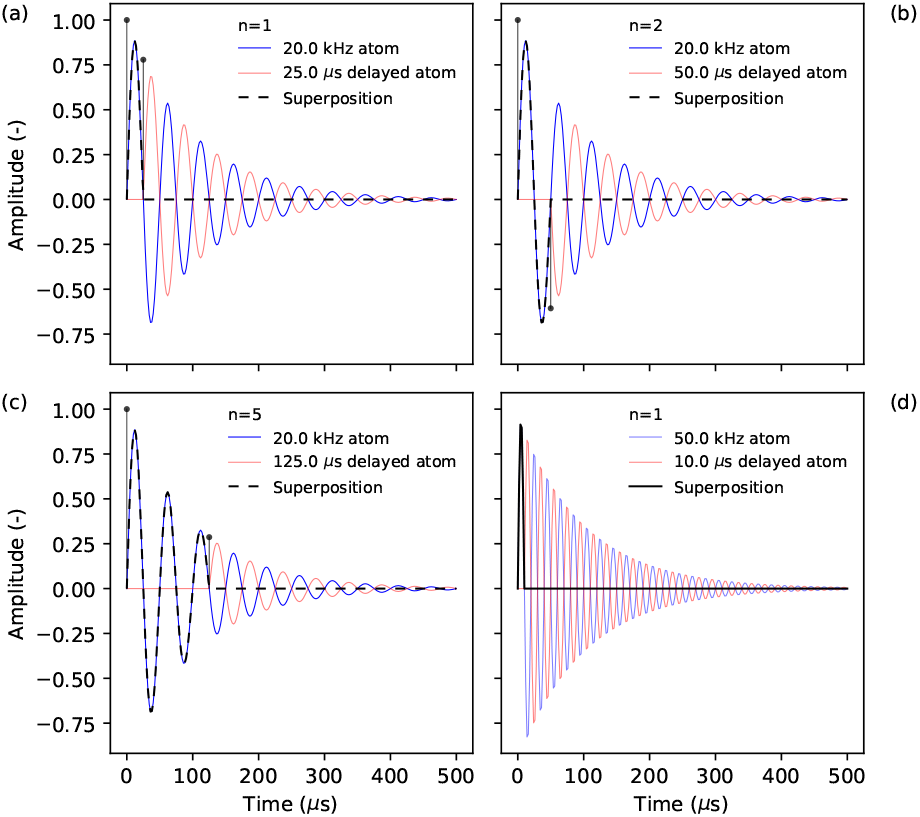
Illustration of temporal localisation achieved through constructive and destructive interference between delayed modal atoms. The black stems correspond to the timing and amplitude of each oscillation. (a) A delayed atom (red) shifted by half a period interferes constructively with the original atom (blue), resulting in a transient with a duration approximately equal to half a period (black). (b) and (c) Delays of two and five periods, respectively, generate progressively longer oscillatory wave packets while maintaining the same underlying resonance. (d) Increasing the resonant frequency reduces the achievable pulse duration for the same relative delay. These examples demonstrate how temporal localisation is controlled by the delay parameter rather than by increasing the damping of the resonant pole.

A dictionary of localised atoms is constructed for each retained resonant frequency. The delay parameter is initially sampled on a uniform grid, after which nearby delay locations are adjusted to coincide with integer multiples of half the modal period. This alignment improves the ability of neighbouring atoms to interfere constructively or destructively, allowing highly localised transient features to be represented with fewer atoms. The resulting collection of delayed resonant atoms forms an overcomplete dictionary, meaning that many different combinations of atoms can represent the same signal. Orthogonal Matching Pursuit (OMP) (Pati et al., 1993) is therefore used to identify a small subset of atoms that accurately reconstructs the transient while minimising the number of active coefficients. The resulting sparse representation separates the resonant structure of the transient from its temporal localisation, yielding a compact and interpretable description of the waveform.

Although the dictionary is sufficiently rich to reconstruct the transient almost exactly (typically correlation coefficients of 0.99–1.00), the objective of the present work is not perfect signal reconstruction. Instead, the number of selected atoms is intentionally limited to obtain a sparse representation that captures the dominant resonant structure while suppressing minor signal variations and noise. This sparse representation facilitates comparison of successive echolocation transients by emphasising repeatable modal features rather than achieving perfect reconstruction. The proposed methodology therefore combines Burg pole estimation (Burg, 1975), sparse representations in over-complete dictionaries (Mallat and Zhang, 1993), and Orthogonal Matching Pursuit (Pati et al., 1993) to obtain a compact modal description of transient signals.

The sparse representation is evaluated using two complementary measures: the reconstruction correlation coefficient and the compression factor,

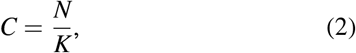

where *N* is the number of samples in the analysis window, and *K* is the number of active atoms selected by OMP.

The correlation coefficient measures how closely the transient is reconstructed, while the compression factor quantifies the compactness of the representation by comparing the number of signal samples in the analysis window to the number of active atoms selected by OMP. A useful decomposition achieves a high reconstruction correlation while maintaining a large compression factor. Increasing the number of active atoms generally improves reconstruction accuracy and honours more click-to-click variability, rather than just the dominant resonant structure. In the present work, the maximum number of active atoms was specified a priori and selected to provide a favourable compromise between reconstruction fidelity and sparsity.

The atoms selected by OMP were subsequently grouped according to their parent resonant pole to obtain the modal decomposition. Summing the delayed atoms associated with each pole yields a single modal response that represents the temporal contribution of that resonance to the transient.

The resonant frequency, quality factor, and amplitude of each modal response were subsequently refined using nonlinear least-squares fitting of a localised damped-resonance. Because the localisation function modifies the waveform just after the onset, this region was excluded from the fit using a temporal mask. The refinement simultaneously provides updated parameter estimates and an uncertainty measure. Refined parameters were accepted only when the relative standard error of the fitted quality factor was below a prescribed threshold; otherwise, the original Burg estimates were retained.

Following parameter refinement, the reconstructed signal is formed from the subset of modal responses that satisfy the confidence criterion. This final quality-control step suppresses poorly constrained modes while retaining the dominant resonant structure of the transient.

## 4 DATASET

The proposed method is evaluated using high-resolution passive acoustic recordings released by MARBEC as part of the DOLPHINFREE project (Lehnhoff et al., 2022, 2025). The dataset contains recordings of free-ranging common dolphins (*Delphinus delphis*) collected in the Northern Bay of Biscay, France. Acoustic measurements were obtained from a single stationary boat using an icListen HF hydrophone with a nominal bandwidth of 10–200 kHz and a sampling frequency of 512 kHz. Observer logs accompanying each recording include information such as dolphin group size, estimated distance and behavioural context. The present study analyses two recordings, each containing several consecutive echolocation transients. Analysing sequences of successive clicks allows the repeatability of the estimated modal parameters to be assessed. The first recording, labelled here as ‘Record 1’, corresponds to the 2020 example described by Lehnhoff et al. (2025) as being randomly selected. The second recording, acquired in 2021, ‘Record 2’, contains a pod that included calves. Although not selected for this purpose, this recording was found to be contaminated by 200 kHz sonar, an engine running at approximately 1200 rpm, and waves, with a period of about two seconds typically associated with swell noise (Elboth et al., 2009). It thus provides a useful opportunity to evaluate the robustness of the proposed method under less favourable recording conditions.

Figure 5 illustrates the quality of ‘Record 1’ containing both echolocation clicks and whistles. Whereas whistles appear as slowly varying tonal components extending over hundreds of milliseconds, the echolocation signals consist of a sequence of short-duration broadband transients. Although the transients dominate the waveform amplitude, their internal temporal structure is only weakly resolved by a conventional spectrogram because of the inherent trade-off between time and frequency resolution. Furthermore, whistle components may overlap individual transients in both time and frequency. Rather than analysing the complete recording, the proposed method processes individual transients in fixed intervals defined by the picked onset and the set window length, allowing each transient to be represented as a sparse combination of damped resonant modes.

**FIGURE 5.**
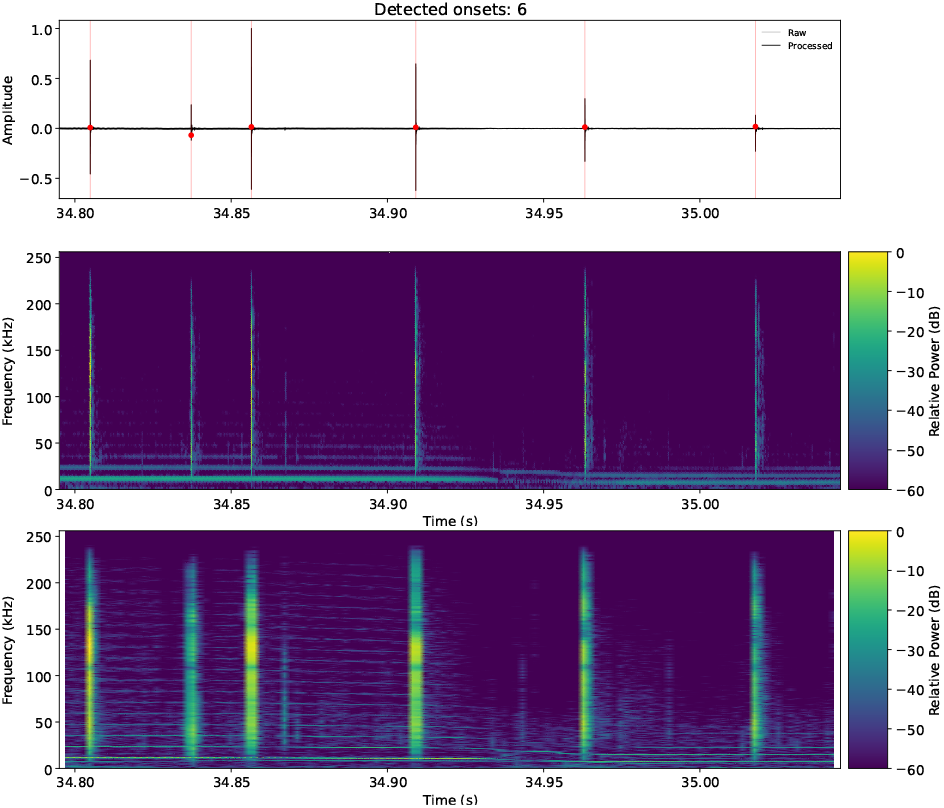
Example recording from the DOLPHINFREE dataset. (Top) Broadband waveform. (Middle) Conventional spectrogram of the same recording optimised for temporal resolution. (Bottom) The spectrogram optimised for frequency resolution.

## 5 RESULTS

In this section, we present the Sparse Modal Representation (SMR) for the two recordings and show the impact of changing the parameterisation. Each plot is made with the same layout as used for the SMR of the synthetic in Figure 1.

### 5.1 Record 1: SMR of individual echolocation clicks

Figure 6 shows the Sparse Modal Representation of six consecutive echolocation clicks from Record 1 spanning a 250 ms interval. Accurate reconstruction is achieved by compressing the data to a ratio of 12.8, requiring only five atoms to represent all resonant modes. The reconstruction quality remains high for all clicks, with maximum normalised cross-correlation coefficients exceeding 0.9. Two distinct click sets can be distinguished. While the temporal activation and amplitudes of the atoms vary between clicks, the resonant frequencies remain remarkably stable, illustrating that the proposed representation captures persistent modal structure while allowing click-to-click variations in temporal localisation.

**FIGURE 6.**
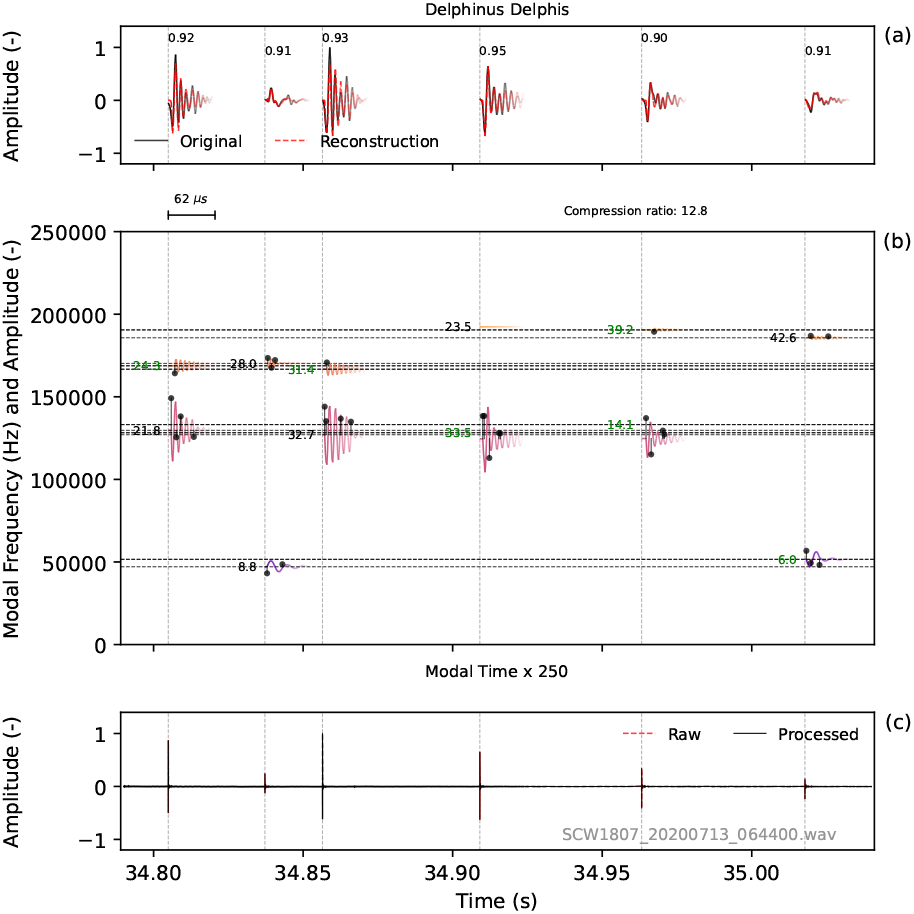
Sparse Modal Representation of six consecutive echolocation clicks from Record 1. The clicks are reconstructed using five atoms distributed over two modes (compression ratio 12.8), yielding correlation coefficients above 0.9.

### 5.2 Reconstruction accuracy versus sparsity

We have demonstrated the construction of a dictionary that is sufficiently rich to almost perfectly match the synthetic transient. Figure 7 shows that this is also the case for the dolphin recording shown in Section 5.1. With the higher number of atoms and an associated drop in compression ratio from 12.8 to 3.6, a perfect reconstruction is achieved. As discussed in Section 3, the objective is not perfect reconstruction but a sparse representation of the dominant resonant structure.

**FIGURE 7.**
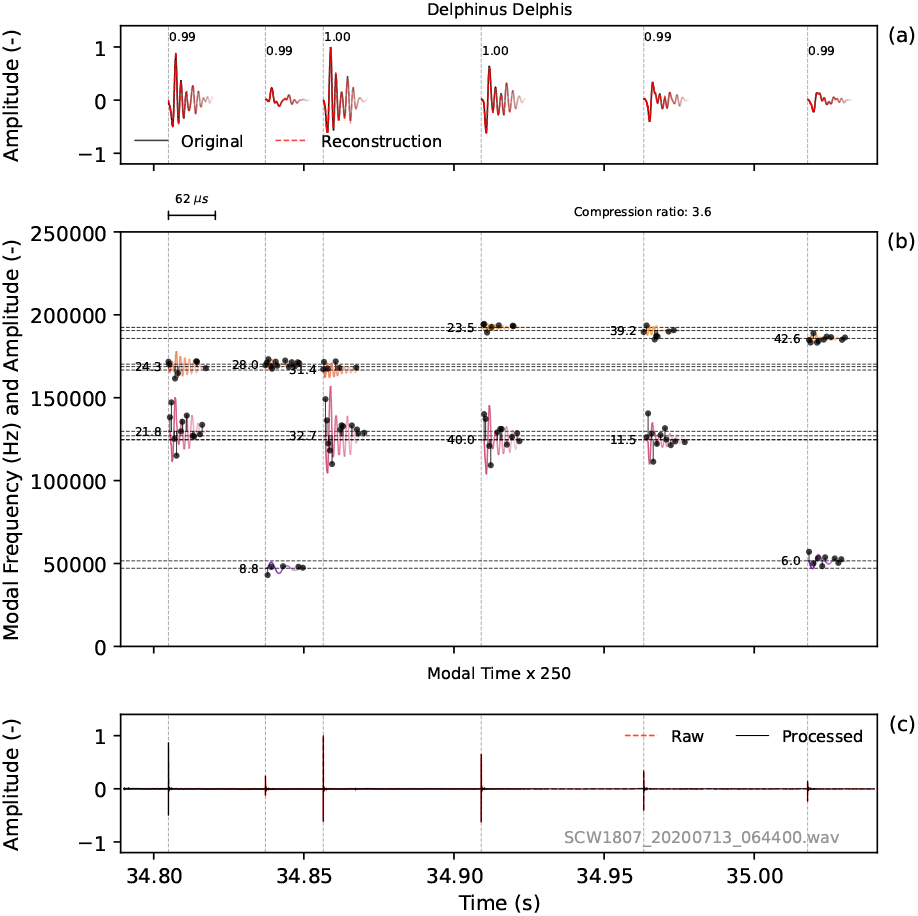
The Sparse Modal Representation for the data in Figure 6 with 18 atoms per mode.

### 5.3 Repeatability of modal parameters

Figure 8 shows a longer section from ‘Record 1’ with 16 consecutive clicks. If interpreted as a single-click sequence, the inter-click intervals alternate between short and long. The difference in modal decomposition makes it more likely that these are two interleaved sequences. The sparse modal representation thus reveals repeated modal patterns across successive clicks that are considerably less apparent in conventional spectrograms. The shift is too large for a Doppler shift, but given the strong directivity of dolphins, this could be explained by a side-to-side motion of the dolphin’s head or possibly by the clicks of two individuals.

**FIGURE 8.**
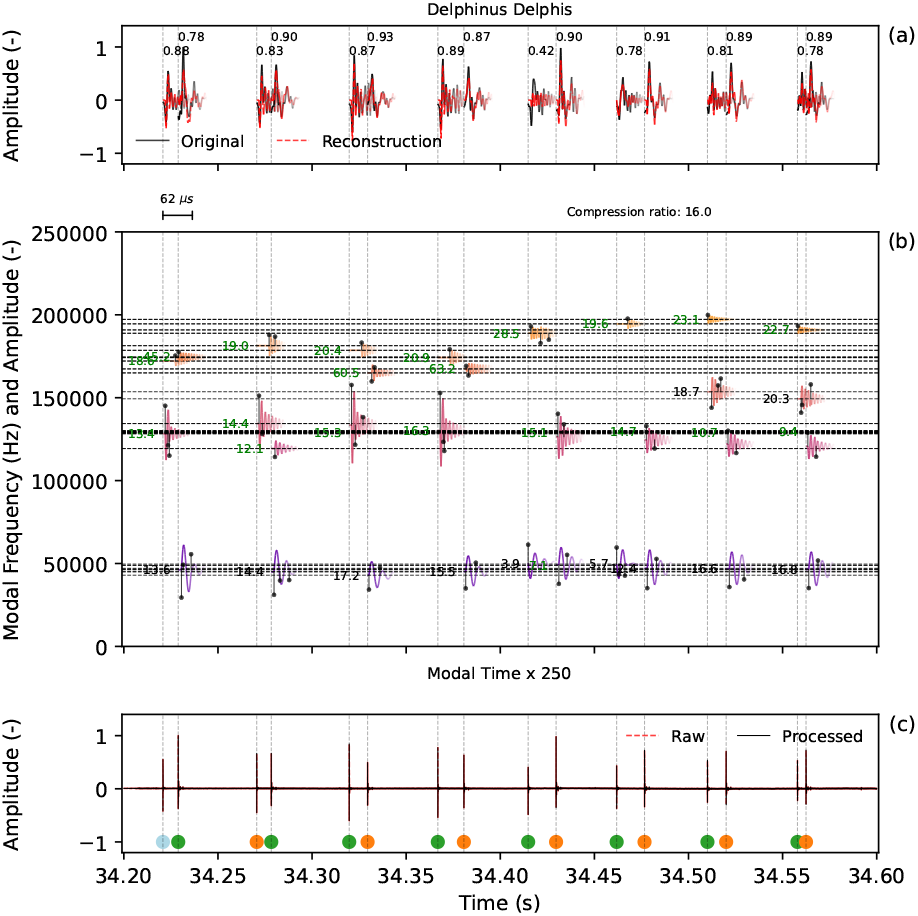
A longer sequence taken from ‘Record 1’ shows alternating short and long intervals. The difference in modal decomposition makes an interleaving of two sequences more likely. Each click is labelled with a dot coloured by cluster using HCA.

**FIGURE 9.**
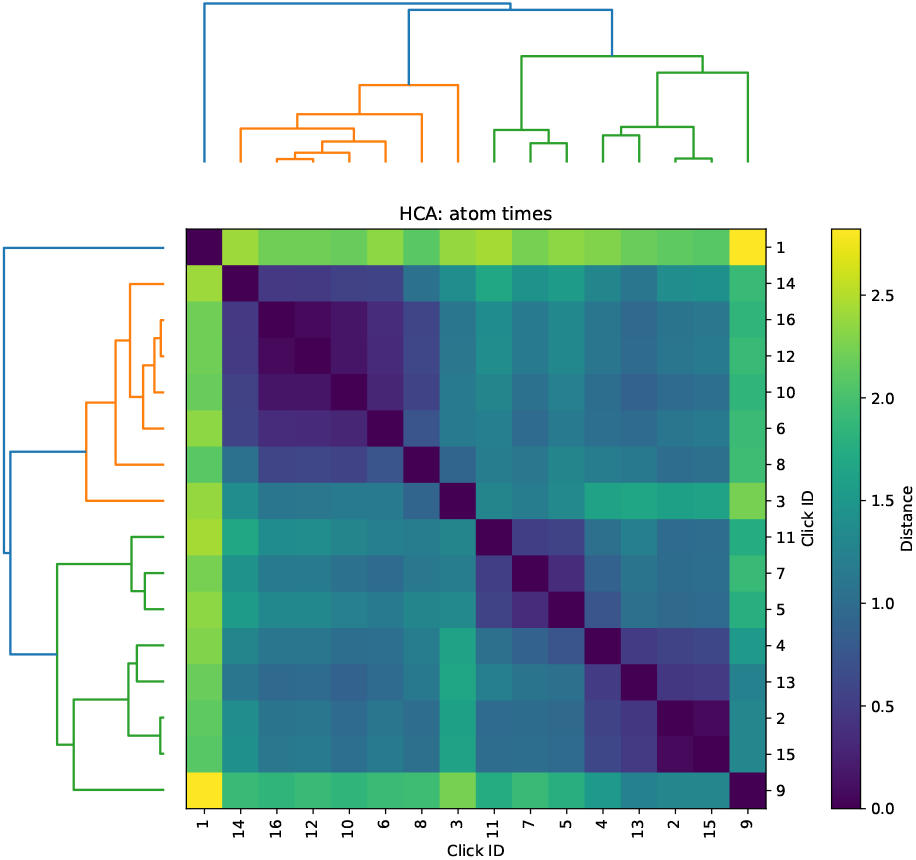
The Hierarchical Cluster Analysis based on atom times. Each click in Figure 8c is labelled by cluster using a coloured dot.

### 5.4 Robustness under noisy recording conditions

The waveform and spectrogram in Figure 10 show that ‘Record 2’ is recorded under noisy conditions. The low-frequency waveand engine noise have low amplitudes and don’t impact the modal decomposition. Figure 11c shows the impact of applying a low-cut filter. Figure 11b shows at 200 kHz a mode with four ‘pings’ per second, which is quite standard for a boat sonar. A FIR low-pass filter effectively removes this noise. In Figure 13, the filtered SMR is shown using just two modes and two to three atoms per mode. By comparing the decomposition with and without the noise, we see that the SMD stability improves. Figure 12 shows the relationship between the correlation coefficient and the compression factor for the high-pass (leaving the sonar noise in the data) and band-pass (filtering out the low-frequency noise and the boat sonar). Removing the boat sonar does increase the risk of overfitting as more poles are available to model the dolphin clicks. Just keeping two modes (Figure 13) shows a reasonable correlation coefficient can be achieved with a higher compression ratio.

**FIGURE 10.**
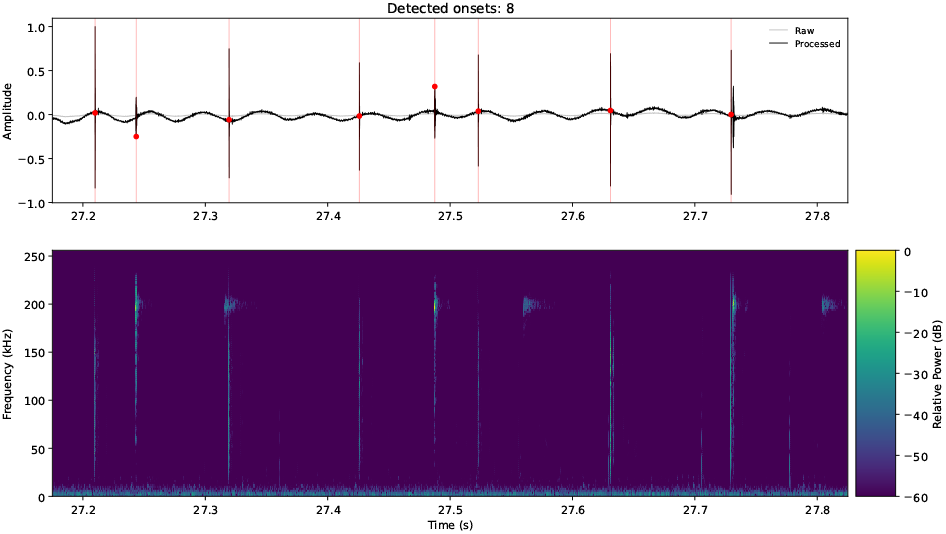
The ‘Record 2’ waveform shows eight high-amplitude clicks (top). The spectrogram (bottom) shows three additional events at 200 kHz, each 100 ms after a high-amplitude event consistent with a boat sonar in a water depth of 75 meters.

**FIGURE 11.**
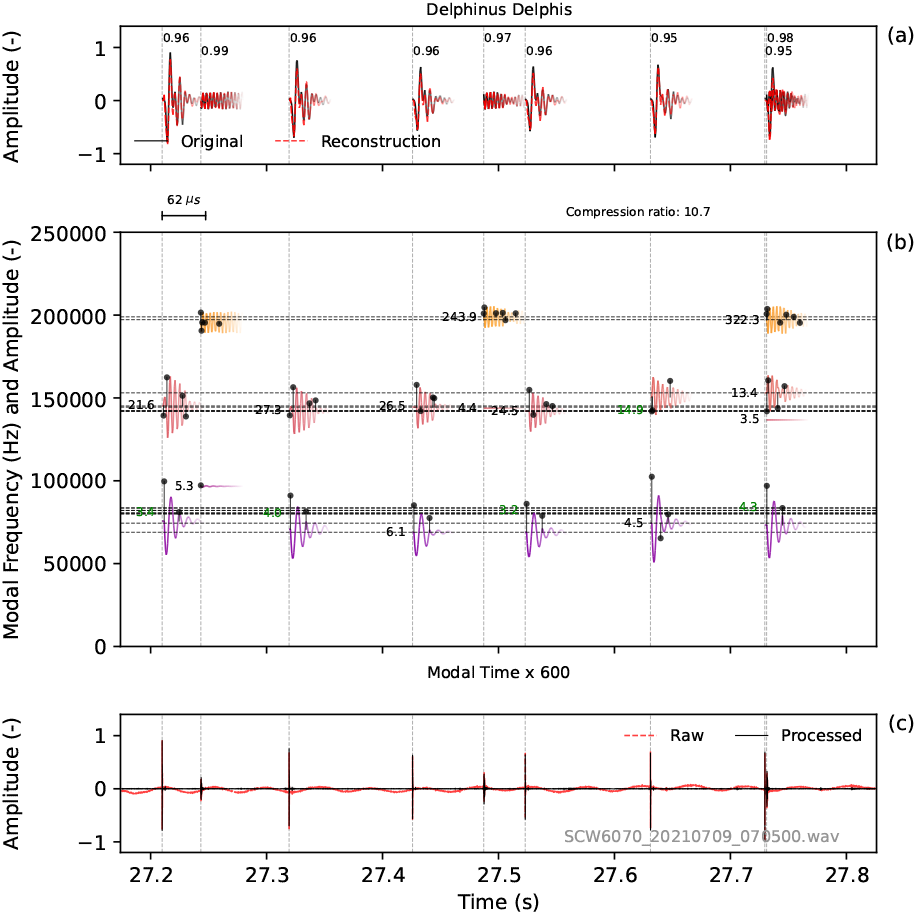
This 600 ms section of ‘Record 2’ shows eight clicks in (c) which are shown with a stretched time scale in (a). The modal decomposition (b) identifies three clicks at 200 kHz associated with a boat sonar.

**FIGURE 12.**
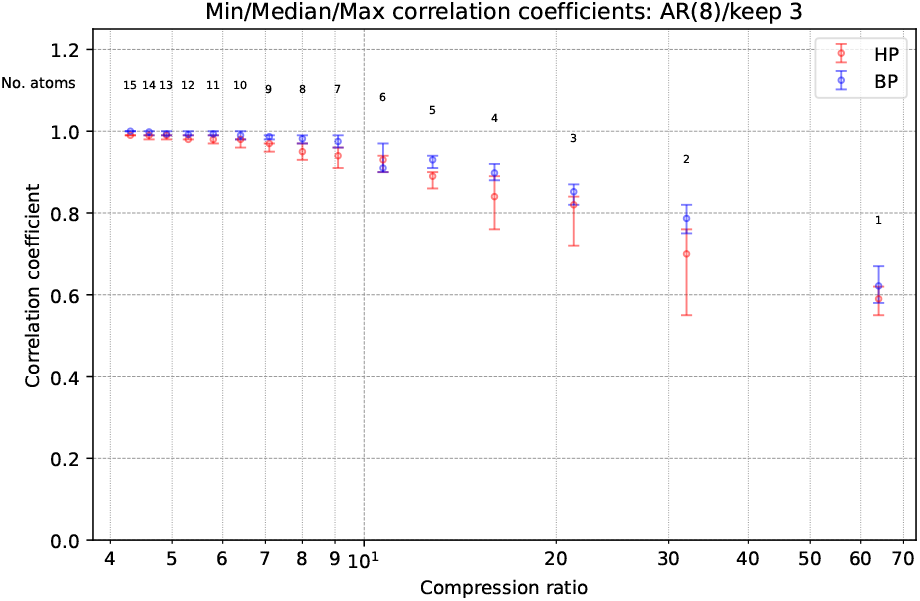
Correlation coefficient vs. Compression factor for Record 2 with a bandpass filter to remove lowand high-frequency noise (BP) and with only the low-frequency noise removed (HP). Each marker shows the minimum, median, and maximum value of the six clicks.

**FIGURE 13.**
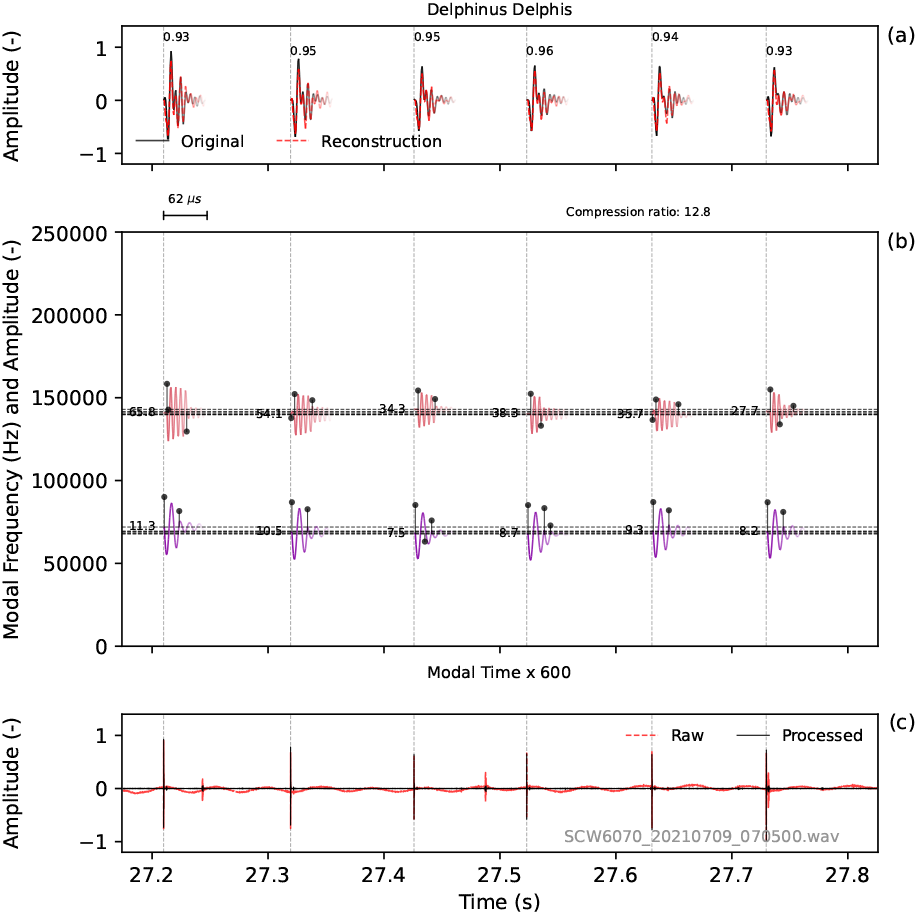
The same section as in Figure 11 after the application of a high-cut filter. A stable decomposition is achieved with a higher compression ratio.

Across both recordings, the proposed sparse modal representation consistently identified a small number of dominant resonant modes. These modes were recovered with high reconstruction accuracy while requiring only a small subset of the available dictionary atoms. Consecutive echolocation clicks exhibited repeatable modal frequencies and quality factors despite variations in temporal activation and recording conditions, suggesting that the representation captures stable resonant characteristics of the emitted transients.

## 6 DISCUSSION

The proposed Sparse Modal Representation (SMR) describes dolphin echolocation transients in terms of a small number of damped resonant modes. Rather than representing each click as a broadband impulse or a collection of stationary frequency components, the method identifies the resonant responses that contribute to the transient together with their temporal activations. The resulting representation is both compact and readily comparable between successive clicks.

Across the analysed recordings, high reconstruction quality was achieved using relatively few active atoms. Correlation coefficients exceeding 0.9 were obtained while maintaining compression factors of approximately 13. Increasing the number of selected atoms allowed almost perfect reconstruction, confirming that the dictionary is sufficiently expressive to reproduce the recorded waveform. However, these experiments also illustrate that maximal reconstruction accuracy is not the primary objective of the proposed method. Restricting the number of active atoms produces a sparser representation that emphasises the dominant resonant structure while suppressing click-to-click variability and measurement noise.

A notable feature of the representation is the stability of the recovered resonant frequencies. In the highest-quality recordings, successive clicks consistently produced clusters of modal frequencies despite variations in click amplitude and temporal localisation. Such stability is difficult to recognise in conventional spectrograms, where the clicks appear primarily as broadband vertical events, but becomes immediately apparent in the sparse modal representation.

The sparse modal representation differs fundamentally from conventional time-frequency analyses. A Fourier transform identifies which frequencies are present, while a wavelet transform identifies which scaled wavelets best represent the signal. In contrast, the proposed representation asks which resonant systems are present and when they are excited. The dictionary is therefore constructed from physically meaningful resonant responses rather than generic mathematical basis functions. This interpretation also remains compatible with existing physical models of dolphin click production, including hypotheses based on internal reflections, while avoiding the need to prescribe a specific source waveform such as a Gabor pulse or Ricker wavelet.

A weighted implementation of Burg’s algorithm was adopted to improve pole estimation for transient signals. Although this modification reduced the number of atoms required to obtain physically meaningful quality factors, a systematic evaluation of transient-aware Burg estimators lies beyond the scope of the present study.

The confidence assessment provides an additional quality-control step. Approximately 30% of the recovered modal responses were rejected based on their parameter uncertainty, yet removing these modes hardly changes the reconstructed waveform. This indicates that OMP naturally assigns negligible coefficients to modes that contribute little to the reconstruction, while the subsequent confidence assessment removes these weakly constrained modes without compromising reconstruction fidelity. Consequently, the reported modal parameters represent only those resonances that are both significant and reliably estimated.

The robustness of the proposed approach was further demonstrated using recordings contaminated by wave noise, boat noise, and boat sonar. The refinement procedure consistently isolated a resonance near 200 kHz that was subsequently identified as a periodic sonar signal with a repetition interval of approximately 4 Hz and a duration of 0.5 ms. This demonstrates that persistent non-biological acoustic sources can be distinguished from dolphin biosonar within the same sparse modal framework.

The proposed Sparse Modal Representation is intended for impulsive, resonance-dominated signals such as odontocete echolocation clicks. Its underlying assumptions of a distinct excitation followed by resonant decay are less appropriate for sustained or frequency-modulated vocalisations, for which alternative time-frequency representations may be more suitable.

The present study has focused on demonstrating the feasibility and interpretability of the sparse modal representation. Future work should investigate larger datasets to determine the biological significance of persistent and intermittent modal resonances, quantify the repeatability of modal parameters across individuals and behavioural contexts, and evaluate whether modal changes provide complementary information for species identification, behavioural classification or studies of biosonar production mechanisms.

## 7 CONCLUSIONS

A Sparse Modal Representation (SMR) has been introduced for analysing dolphin echolocation transients. The proposed method combines Burg autoregressive pole estimation with a sparse dictionary of localised damped resonant atoms and Orthogonal Matching Pursuit to create a compact representation. This representation simultaneously estimates resonant frequency, damping, temporal localisation, and reconstruction quality.

When applied to high-resolution recordings of free-ranging common dolphins, the method demonstrated that individual clicks can be accurately represented using a small number of modal atoms. This approach maintains high reconstruction correlation and substantial compression. The resulting sparse modal representation reveals stable clusters of resonant frequencies across successive clicks. This clustering naturally separates persistent resonant structures from weaker or poorly constrained components through confidence-based parameter refinement.

Unlike conventional time-frequency representations, the proposed approach characterises a transient in terms of its underlying resonant systems and their temporal activations. This provides an interpretable description of click structure, facilitating comparisons between successive echolocation transients. It also remains compatible with existing physical interpretations of dolphin click production.

While demonstrated here for dolphin biosonar, this methodology can be applied to a wider range of transient signals. The Sparse Modal Representation offers not only an efficient representation of signals but also a resonance-centred framework for interpreting transient acoustic signals.

## 8 DATA AVAILABILITY

The recordings analysed in this study are part of the publicly available DOLPHINFREE dataset described by Lehnhoff et al. (2025). The dataset is distributed under the Creative Commons Attribution 4.0 International (CC BY 4.0) License.

## 9 ACKNOWELDGEMENTS

The encouragement of Ben Hillier and the insightful comments of Sarab Sethi are gratefully acknowledged.

## APPENDIX A SPARSE MODAL DECOMPOSITION

Consider a sampled transient signal

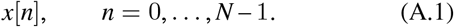

The objective is to represent the transient as a sparse combination of localized damped modal atoms.

### A.1 Autoregressive Pole Estimation Using Burg’s Method

An autoregressive model of order *p* is fitted to the signal,

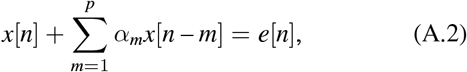

where *α*_*m*_ are the autoregressive coefficients and *e*[*n*] is the prediction error.

The associated autoregressive polynomial is

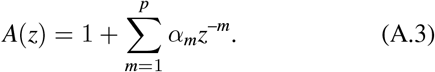

Burg’s method estimates the coefficients *α*_*m*_ by minimizing the combined forward and backward prediction errors.

#### A.2 Pole Extraction

The discrete-time poles are obtained as the roots of

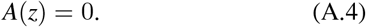

Let

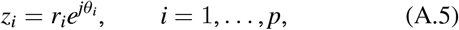

denote the resulting poles.

Only stable poles satisfying

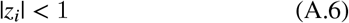

are retained, thereby excluding exponentially growing modes.

For real-valued signals, complex poles occur in conjugate pairs 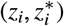. Since the dictionary atoms are constructed from the real part of the modal response, only the poles with positive imaginary part are retained. The corresponding conjugate poles are implicitly represented through

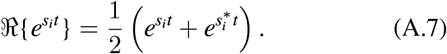

The *N*_keep_ least damped resonant poles are selected by sorting the stable poles according to descending radius |*z*_*i*_|, corresponding to poles closest to the unit circle.

#### A.3 Continuous-Time Pole Representation

Using the sampling interval

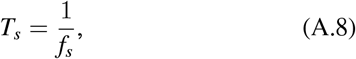

the discrete-time poles are mapped to continuous-time poles,

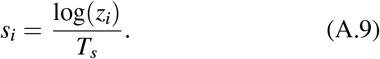

Writing

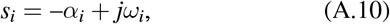

the modal frequency becomes

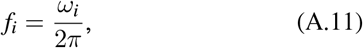

and the modal quality factor is

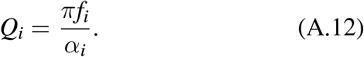

#### A.4 Localized Modal Atoms

For each pole *s*_*i*_ and delay location *τ*_*j*_, a localized modal atom is defined as

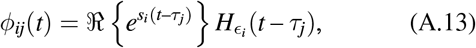

where

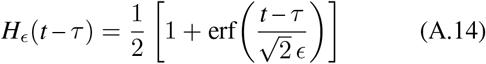

is a smooth approximation of the Heaviside step function. The parameter *ϵ* controls the transition width and therefore the temporal localization of the atom.

The localization width is chosen as

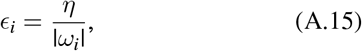

where *η* is a user-defined localization parameter.

The delayed atoms provide localised activation of the resonant modes. Through constructive and destructive interference between delayed atoms, finite-duration wave packets can be synthesized from lightly damped resonances.

#### A.5 Dictionary Construction

Given the candidate delay locations

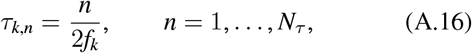

formed from half-period increments of each selected resonant pole, and the uniform reference grid

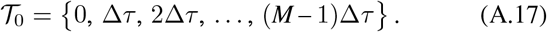

The delay grid is adapted according to

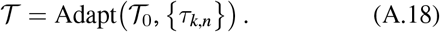

The operator Adapt(*·*) replaces the nearest locations in the uniform grid *T*_0_ by the modal delay locations *τ*_*k,n*_, removes duplicate values, and sorts the resulting delay set in ascending order.

The resulting adapted delay set contains

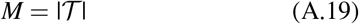

delay locations.

A dictionary matrix is formed by concatenating all localized modal atoms,

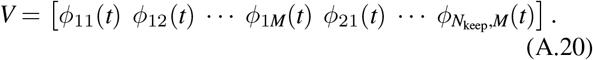

The total number of dictionary atoms is

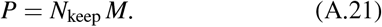

#### A.6 Sparse Coefficient Estimation

Given the dictionary matrix *V*, a sparse coefficient vector

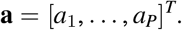

is determined using Orthogonal Matching Pursuit (OMP) by solving

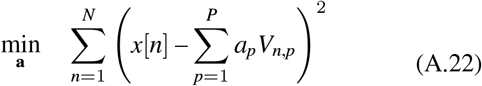

where

- *x*[*n*] is the signal sample at time index *n*,
- *V*_*n,p*_ is the *n*-th sample of the *p*-th dictionary atom,
- *a*_*p*_ is the corresponding sparse coefficient,
- *P* is the total number of dictionary atoms, and
- *∥***a***∥*_0_ denotes the number of non-zero coefficients in **a**.

The optimization seeks the sparsest representation of the signal using at most *N*_atoms_ active dictionary atoms while minimizing the squared reconstruction error.

The resulting sparse coefficient vector identifies the subset of dictionary atoms that best reconstructs the signal while limiting model complexity.

#### A.7 Grouped Modal Responses

Let *a*_*ij*_ denote the sparse coefficient associated with pole *s*_*i*_ and delay *τ*_*j*_ such that

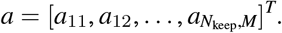

The grouped response associated with pole *s*_*i*_ is defined as

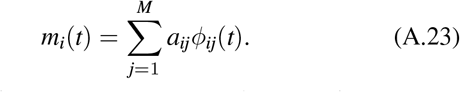

Each grouped response represents the contribution of a single resonant pole across all active delay locations.

#### A.8 Parameter Refinement

The grouped modal responses provide isolated resonant components from which the modal parameters may be refined.

For each grouped response, the model

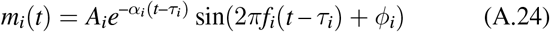

is fitted by nonlinear least squares to the fully activated portion of the signal, excluding the initial localization interval. The refined modal parameters comprise the modal frequency *f*_*i*_, quality factor

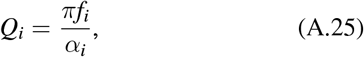

the modal amplitude *A*_*i*_, and the phase *ϕ*_*i*_.

#### A.9 Confidence Assessment

The covariance matrix returned by the nonlinear least-squares fit is used to assess the reliability of the refined modal parameters.

For each fitted parameter,

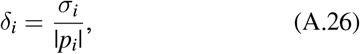

where *σ*_*i*_ denotes the estimated standard deviation obtained from the covariance matrix and *p*_*i*_ the corresponding fitted parameter.

Refined parameters are accepted only when the relative uncertainty falls below a prescribed threshold and remain consistent with the initial Burg estimate. Otherwise the original Burg estimate is retained.

#### A.10 Signal Reconstruction

The reconstructed signal is obtained as the sum of all grouped modal responses,

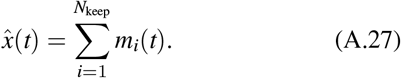

Equivalently,

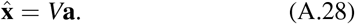

#### A.11 Compression Ratio

The number of active atoms is defined as

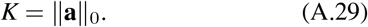

For a signal containing *N* samples, the compression ratio is defined as:

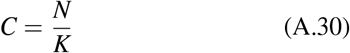

Larger values of *C* indicate that the transient can be represented using fewer active modal atoms.

The sparse modal decomposition provides a continuous transition between a parametric representation (*K ≪ N*) and a non-parametric representation (*K → N*), depending on the number of active dictionary atoms retained.

#### A.12 Summary

The complete decomposition procedure may be summarized as

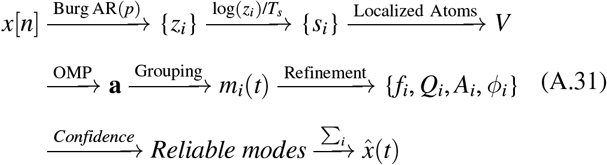

## APPENDIX B SPARSE MODAL REPRESENTATION: EXAMPLES

### B.1 Toothed Whales

Figure B.1 shows six echolocation clicks part of a Sperm Whale recording, provided by Mahmudov (2025) under a CC0 1.0 license as a Kaggle dataset. Noteworthy is the stable low-frequency tail of these echolocation clicks. The mode at approximately 7500 Hz is very stable. The highest frequency mode is variable.

**FIGURE B.1.**
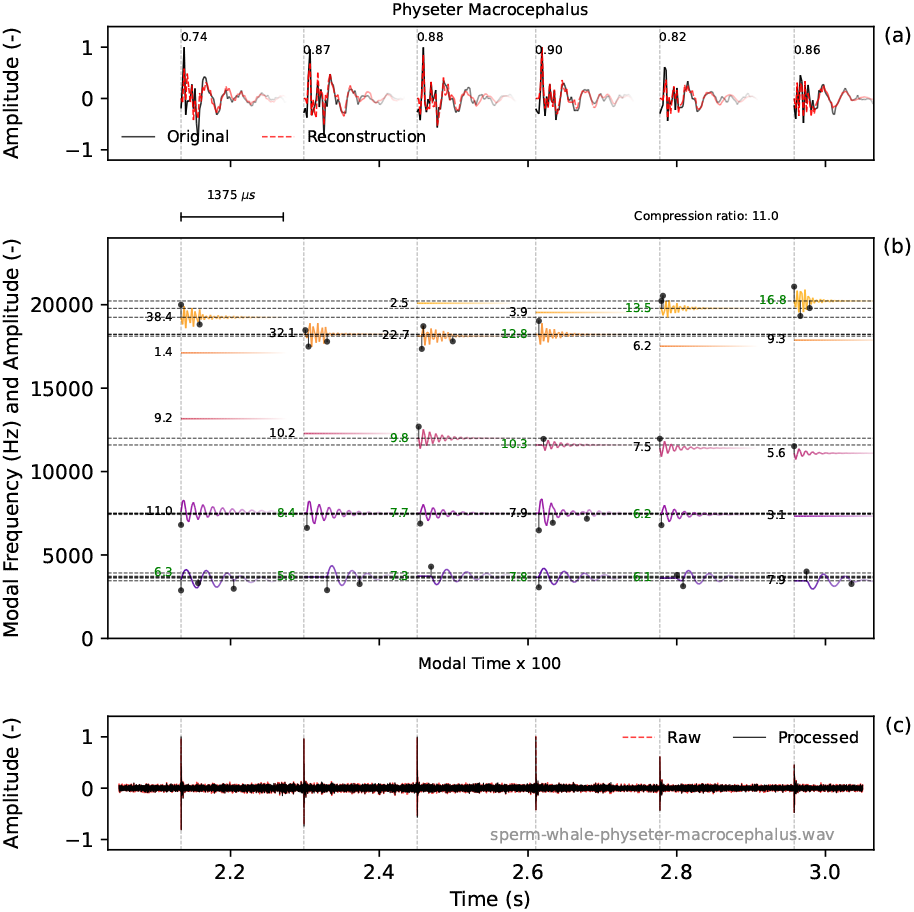
Sparse Modal Representation of a Sperm whale.

The clicks in Figure B.2 slowly decay over approximately 100 ms. This long tail could, for example, consist of many reflections. The decomposition of the first three milliseconds, although consisting of several bursts, is highly repeatable.

**FIGURE B.2.**
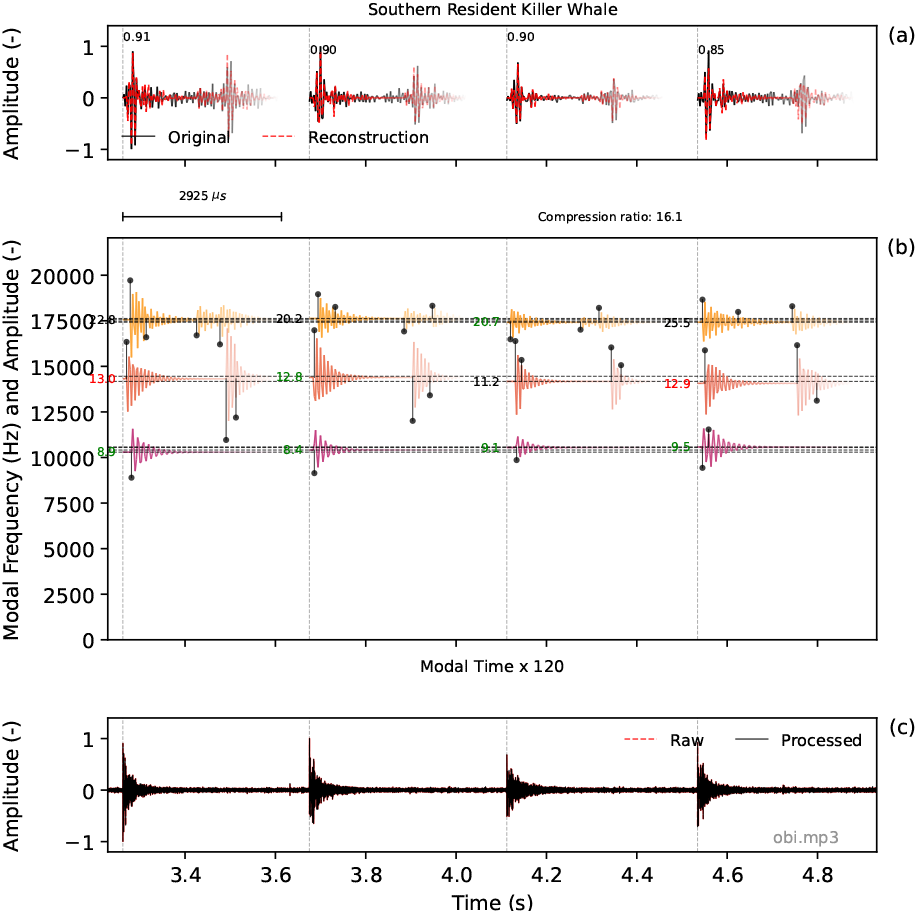
Sparse Modal Representation of a Southern Resident Killer Whale.

#### B.2 Bats

Here these specific bats fit into the categories of acoustic mechanism and signal design:

- Egyptian Fruit Bat: Tongue-Clicking (Lingual) Echolocation: They do not use their voice boxes for sonar. Instead, they snap their tongues against the inside of their mouths to produce audible clicking sounds that help them navigate pitch-black caves. Figure B.3 shows the SMR of an Egyptian Fruit bat. The data is an example from the Nature Science dataset *‘An annotated dataset of Egyptian fruit bat vocalizations’* (Prat et al., 2017).
- Alcathoe Bat (*Myotis alcathoe*): Laryngeal Echolocation / Mouth Emitter / Frequency Modulated (FM) This tiny woodland bat shouts ultrasonic calls through its mouth. It utilizes steep, broadband FM sweeps that drop from around 120 kHz down to 40 kHz (Jan et al., 2010). The SMD in Figure B.4 shows a frequency drop from 121 kHz to 52.6 kHz. Figure B.5 shows these frequencies plotted as lines on the spectrogram analysis. The frequencies found by the spectrogram analysis are somewhat lower. Audio source: PeerSonic webpage (PeerSonic, 2026). Data provided as ‘Open Source’.
- Noctule Bat (*Nyctalus noctula*): Laryngeal Echolocation / Mouth Emitter / Mixed (FM-CF) Noctule Bat calls range from 20 to 45kHz with a peak at 25kHz. A characteristic ‘chip chop’ with occasional clicks is heard during feeding on a bat detector. The echolocation click is a mix of ‘frequency modulated’ and ‘constant frequency’ (or Quasi-FM). A subtle frequency-modulated onset and alternating higher and lower frequencies are visible in Figure B.6.

While a single exponential damping model would struggle to represent a frequency-modulated click, we can break it down into very short atoms. Figure B.7 demonstrates that for the click at 0.96 seconds, the SMD successfully resolves a frequency drop of approximately 8 kHz.

**FIGURE B.3.**
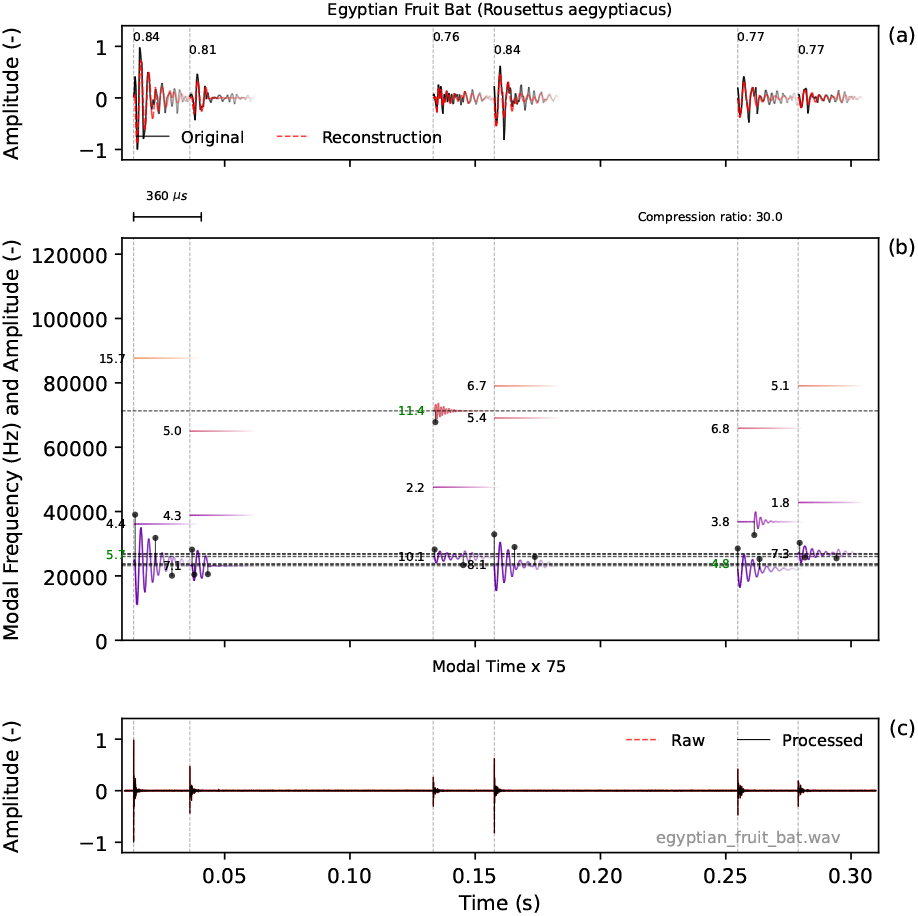
Sparse Modal Representation of an Egyptian Fruit bat.

**FIGURE B.4.**
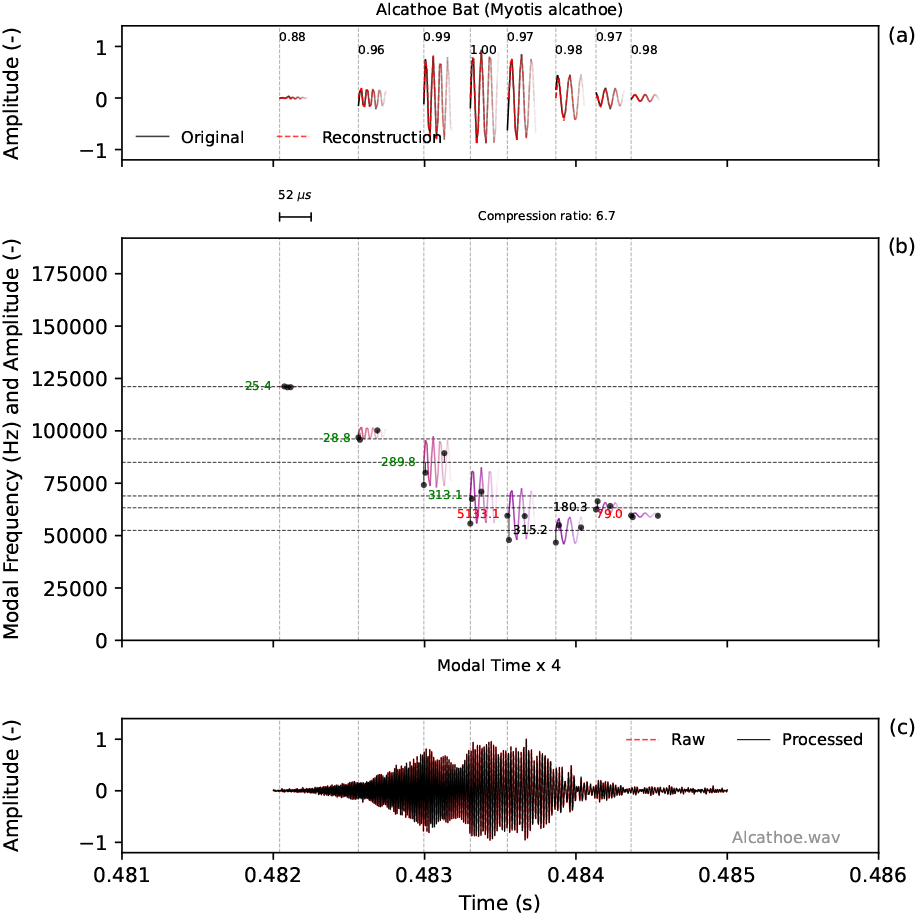
Sparse Modal Representation of an Alcathoe bat.

**FIGURE B.5.**
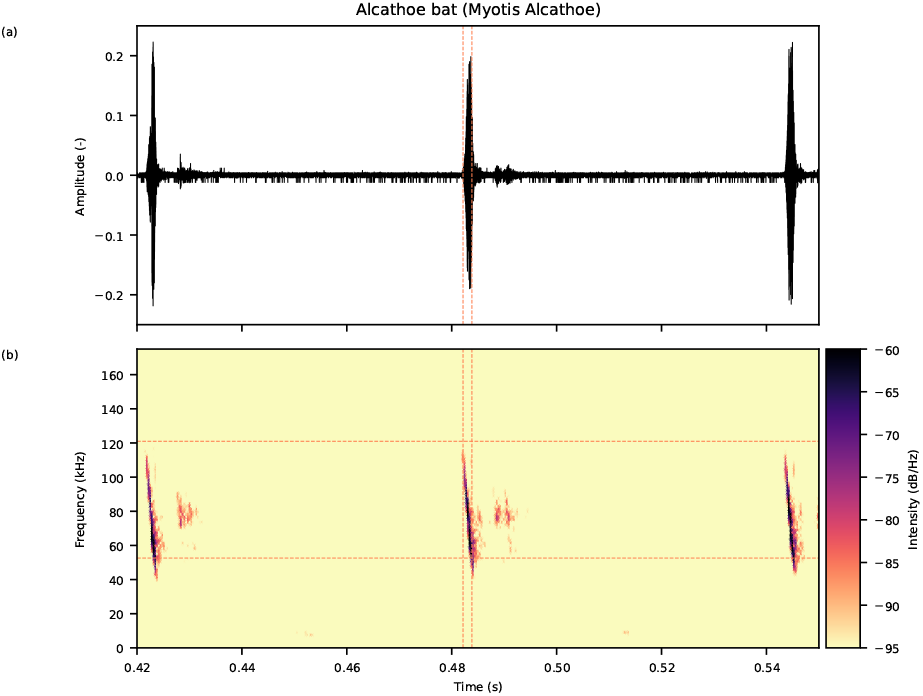
Spectrogram of Alcathoe bat echolocation clicks.

Figure B.8 provides a zoomed-in spectrogram of the click at 2.5 seconds in Figure B.6. The spectrogram’s maxima can be used to derive a frequency versus time curve (orange dashed line) or a ‘Fourier-based estimate’. Alternatively, we can observe the frequency drift relative to the Burg-estimated frequency (black dashed line) to estimate the instantaneous frequency.

To minimise numerical noise in direct frequency differentiation, the instantaneous frequency drift is estimated by fitting a smooth integrated phase profile of a hybrid exponential-linear model (the red line). The resulting ‘damping’ and ‘kinematic’ parameters describe a mechanical model that could be used to study passive viscoelastic tissue damping and muscular sweep kinematics within the phase domain.

Audio source: PeerSonic webpage (PeerSonic, 2026). Data provided as ‘Open Source’.

**FIGURE B.6.**
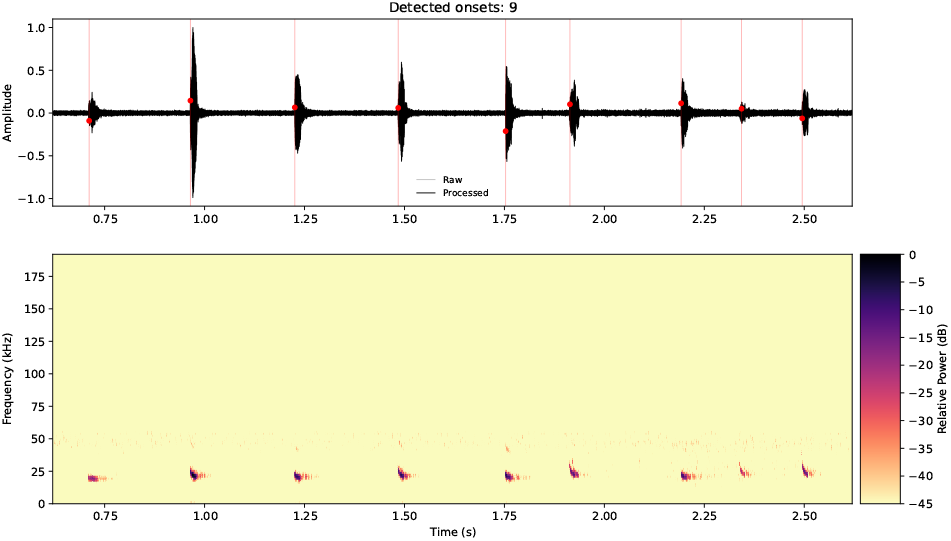
Spectrogram of a series of Noctule Bat echolocation clicks.

**FIGURE B.7.**
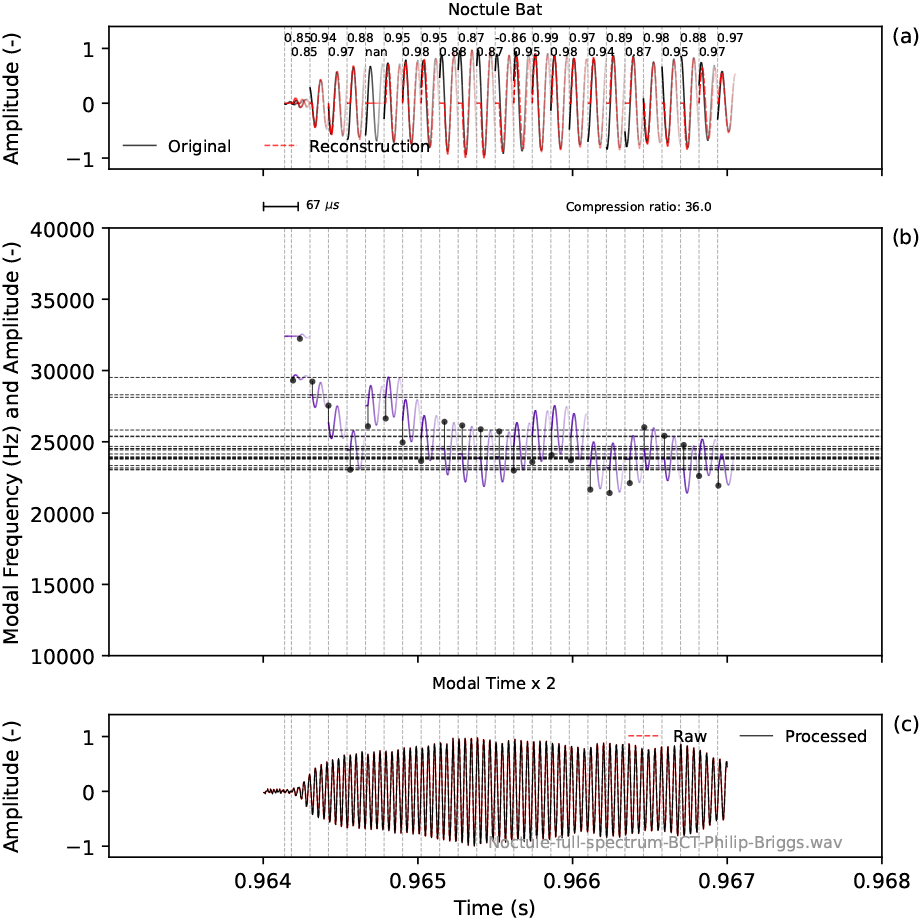
Detail of Noctule SMR click shown at 0.96 seconds in Figure B.6.

**FIGURE B.8.**
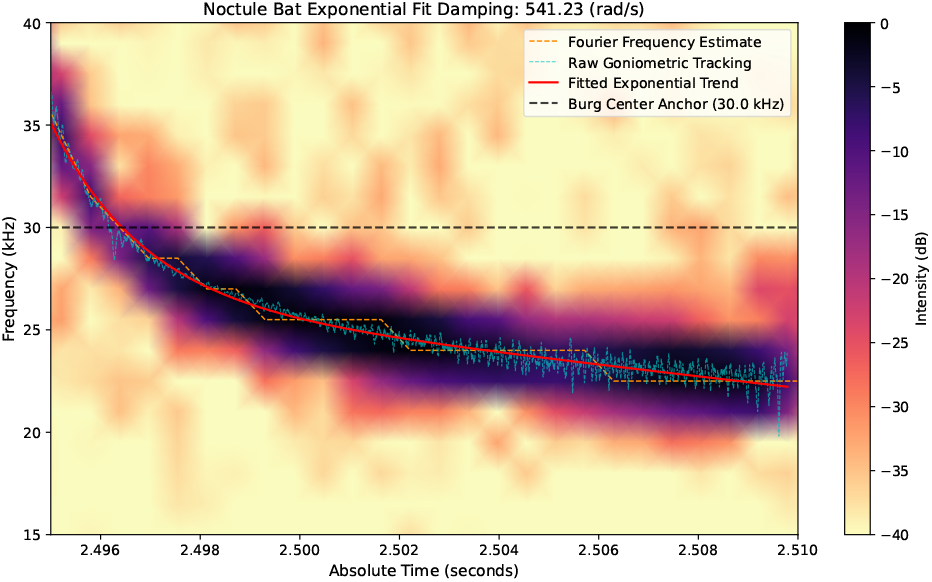
Exponential fit to click at 2.5 seconds in Figure B.6.

#### B.3 Birds

Echolocation has been observed for several bird species: the South American Olibird (*Steatornis caripensis*) and several swiflets of the *Aerodramus* and *Collocalia* genera. The click frequencies fall within the human audible range (Brinkløv et al., 2013).

The SMR of a Mossy-nest Swiftlet in Figure B.9 shows two distinct modes. Applying a bandpass filter improved the stability of the pole frequencies. Audio source: bbc.co.uk – © copyright 2026 BBC (Colbeck, 1986)

Figure B.10 shows that the echolocation clicks are well within the human hearing range. A reasonable representation is achieved with the modes occurring at harmonic intervals of 2.5 kHz. In Figure B.11 much weaker peaks are included in the analysis. The fundamental frequency and harmonics show a slightly higher frequency. Audio source: bbc.co.uk – © copyright 2026 BBC (Bird, 1982)

**FIGURE B.9.**
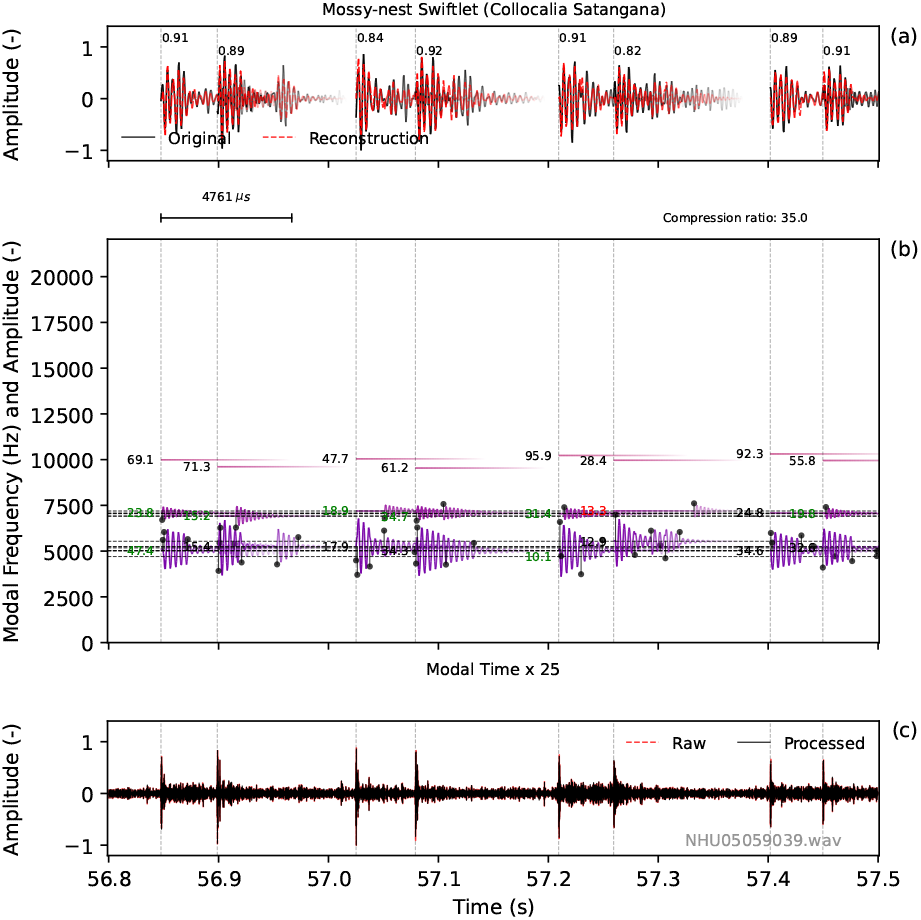
Sparse Modal Representation of a Mossy-nest Swiftlet.

**FIGURE B.10.**
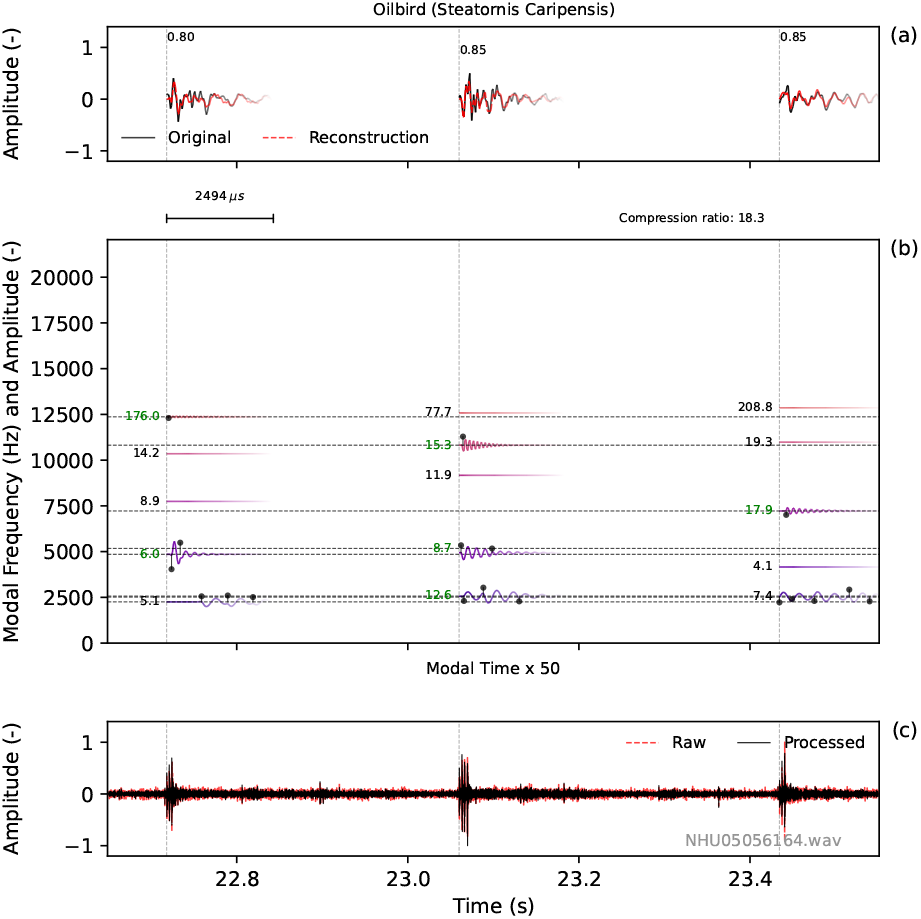
Sparse Modal Representation of an Oilbird.

**FIGURE B.11.**
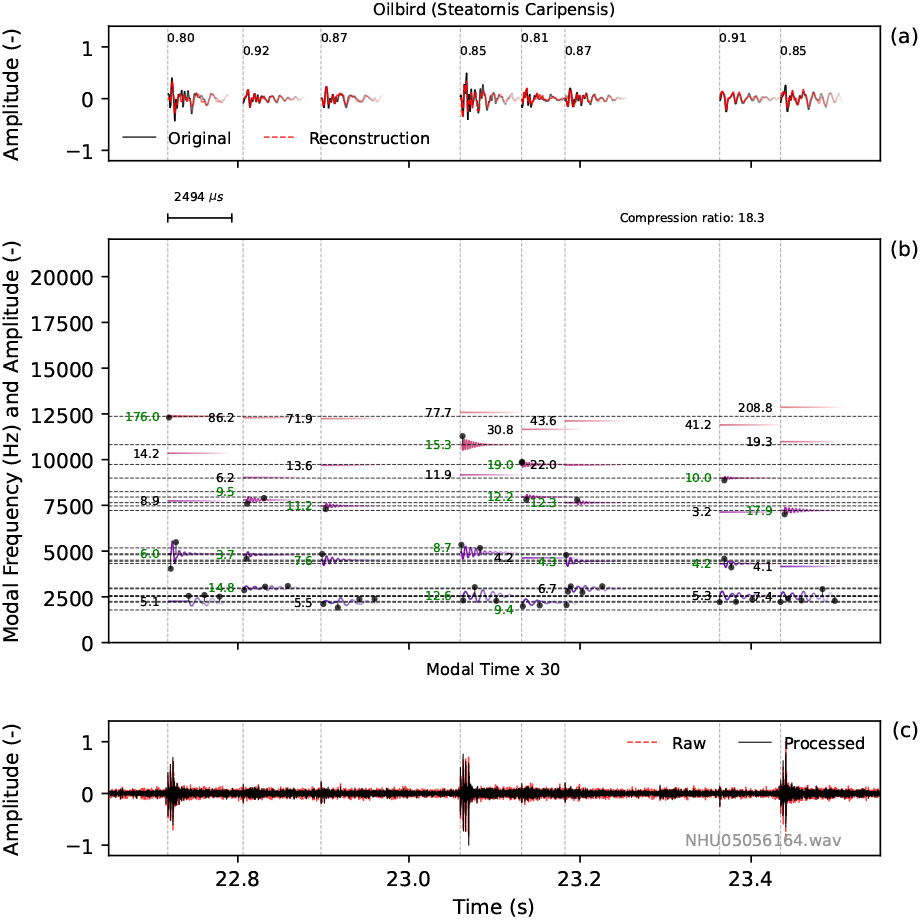
Sparse Modal Representation of one or more Oilbirds.

## REFERENCES

Au, W.W.L. (2015) History of dolphin biosonar research. Acoustics Today, 11, 10–17. URL https://acousticstoday.org/history-of-dolphin-biosonar-research-by-whitlow-w-l-au/

Bird, L. (1982) Bbc sound effects: Oilbird (several birds in cave). URL https://sound-effects.bbcrewind.co.uk/search?q=NHU05056164

Borino, G. & Muscolino, G. (1986) Mode-superposition methods in dynamic analysis of classically and non-classically damped linear systems. Earthquake Engineering & Structural Dynamics, 14, 705 – 717. doi:10.1002/eqe.4290140503.

Brinkløv, S., Fenton, M.B. & Ratcliffe, J.M. (2013) Echolocation in oilbirds and swiftlets. Frontiers in Physiology, 4 MAY. doi:10.3389/fphys.2013.00123.

Burg, J.P. (1975) Maximum entropy spectral analysis. Ph. D. thesis, Stanford University, Department of Geophysics,. URL http://sepwww.stanford.edu/theses/sep06/

Colbeck, M. (1986) Bbc sound effects: Mossy-nest swiftlet. URL https://sound-effects.bbcrewind.co.uk/search?q=NHU05059039

Elboth, T., Reif, B.A.P. & Øyvind Andreassen (2009) Flow and swell noise in marine seismic data. Geophysics, 74, Q17– Q25. doi:10.1190/1.3078403. URL https://doi.org/10.1190/1.3078403

Jan, C.M., Frith, K., Glover, A.M., Butlin, R.K., Scott, C.D., Greenaway, F. et al. (2010) Myotis alcathoe confirmed in the uk from mitochondrial and microsatellite dna. Acta Chiropterologica, 12, 471–483. doi:10.3161/150811010X538043.

Kamminga, C., Stuart, A.C. & Silber, G.K. (1996) Investigations on cetacean sonar xi: Intrinsic comparison of the wave shapes of some members of the phocoenidae family. Aquatic Mammals, 22, 45–55.

Kamminga, C. & Wiersma, H. (1981) Investigations on cetacean sonar ii. acoustical similarities and differences in odontocete sonar signals. Aquatic Mammals, 8, 41–62.

Lehnhoff, L., Glotin, H., Bernard, S., Dabin, W., Gall, Y.L., Menut, E. et al. (2022) Behavioural responses of common dolphins delphinus delphis to a bio-inspired acoustic device for limiting fishery by-catch. Sustainability (Switzerland), 14. doi:10.3390/su142013186.

Lehnhoff, L., Glotin, H., Gall, Y.L., Menut, E., Peltier, H., Spitz, J. et al. (2025) High-resolution acoustic recordings of wild free-ranging short-beaked common dolphins for etho-acoustical and repertoire studies. Earth System Science Data, 17, 4495–4511. doi:10.5194/essd-17-4495-2025.

Leighton, T., Richards, S. & White, P. (2004) Marine mammal signals in bubbly water. Sonar Signal Processing Pt, 26266.

Mahmudov, A. (2025) Noaa fisheries, whale sound records. URL https://www.kaggle.com/datasets/asimmahmudov/whale-sounds-dataset/data

Mallat, S.G. & Zhang, Z. (1993) Matching pursuits with time-frequency dictionaries. IEEE Transactions on Signal Processing, 41, 3397–3415. doi:10.1109/78.258082.

Pati, Y.C., Rezaiifar, R. & Krishnaprasad, P.S. Orthogonal matching pursuit: recursive function approximation with applications to wavelet decomposition. In: Proceedings of 27th Asilomar Conference on Signals, Systems and Computers, 1993, pp. 40–44 vol.1.

PeerSonic (2026) Peersonic samples and sound. URL https://www.peersonic.co.uk/files/116/Sound-Samples/17/Noctule

Prat, Y., Taub, M., Pratt, E. & Yossi, Y. (2017) Example of egyptian fruit bat echolocation. URL https://figshare.com/articles/dataset/Example_of_Egyptian_fruit_bat_echolocation_1_/5015996?backTo=/collections/An_annotated_dataset_of_Egyptian_fruit_bat_vocalizations_across_varying_contexts_and_during_vocal_ontogeny/3666502

Wahlberg, M., Jensen, F.H., Soto, N.A., Beedholm, K., Bejder, L., Oliveira, C. et al. (2011) Source parameters of echolocation clicks from wild bottlenose dolphins ( tursiops aduncus and tursiops truncatus). The Journal of the Acoustical Society of America, 130, 2263–2274. doi:10.1121/1.3624822.

Wei, C., Au, W.W.L., Ketten, D.R. & Zhang, Y. (2018) Finite element simulation of broadband biosonar signal propagation in the near- and far-field of an echolocating atlantic bottlenose dolphin ( tursiops truncatus). The Journal of the Acoustical Society of America, 143, 2611–2620. doi:10.1121/1.5034464.

Wei, C., Erbe, C., Smith, A.B. & Yang, W.C. (2024) Validated 3d finite-element model of the risso’s dolphin (grampus griseus) head anatomy demonstrates gular sound reception and channelling through the mandibular fats. Bioinspiration and Biomimetics, 19. doi:10.1088/1748-3190/ad7344.

